# Euplotid: A quantized geometric model of the eukaryotic cell

**DOI:** 10.1101/170159

**Authors:** Diego Borges-Rivera

## Abstract

1

Life continues to shock and amaze us, reminding us that truth is far stranger than fiction. http://Euplotid.io is a quantized geometric model of the eukaryotic cell, an attempt at quantifying the incredible complexity that gives rise to a living cell by beginning from the smallest unit, a quanta. Starting from the very bottom we are able to build the pieces which when hierarchically and combinatorially combined produce the emergent complex behavior that even a single celled organism can show. Euplotid is composed of a set of quantized geometric 3D building blocks and constantly evolving dockerized bioinformatic pipelines enabling a user to build and interact with the local regulatory architecture of every gene starting from DNA-interactions, chromatin accessibility, and RNA-sequencing. Reads are quantified using the latest computational tools and the results are normalized, quality-checked, and stored. The local regulatory architecture of each gene is built using a Louvain based graph partitioning algorithm parameterized by the chromatin extrusion model and CTCF-CTCF interactions. Cis-Regulatory Elements are defined using chromatin accessibility peaks which are mapped to Transcriptional Start Sites based on inclusion within the same neighborhood. Deep Neural Networks are trained in order to provide a statistical model mimicking transcription factor binding, giving the ability to identify all Transcription Factors within a given chromatin accessibility peak. By in-silico mutating and re-applying the neural network we are able to gauge the impact of a transition mutation on the binding of any transcription factor. The annotated output can be visualized in a variety of 1D, 2D, 3D and 4D ways overlaid with existing bodies of knowledge such as GWAS results or PDB structures. Once a particular CRE of interest has been identified a Base Editor mediated transition mutation can then be performed in a relevant model for further study.

Figure 0.1:
Graphical Abstract

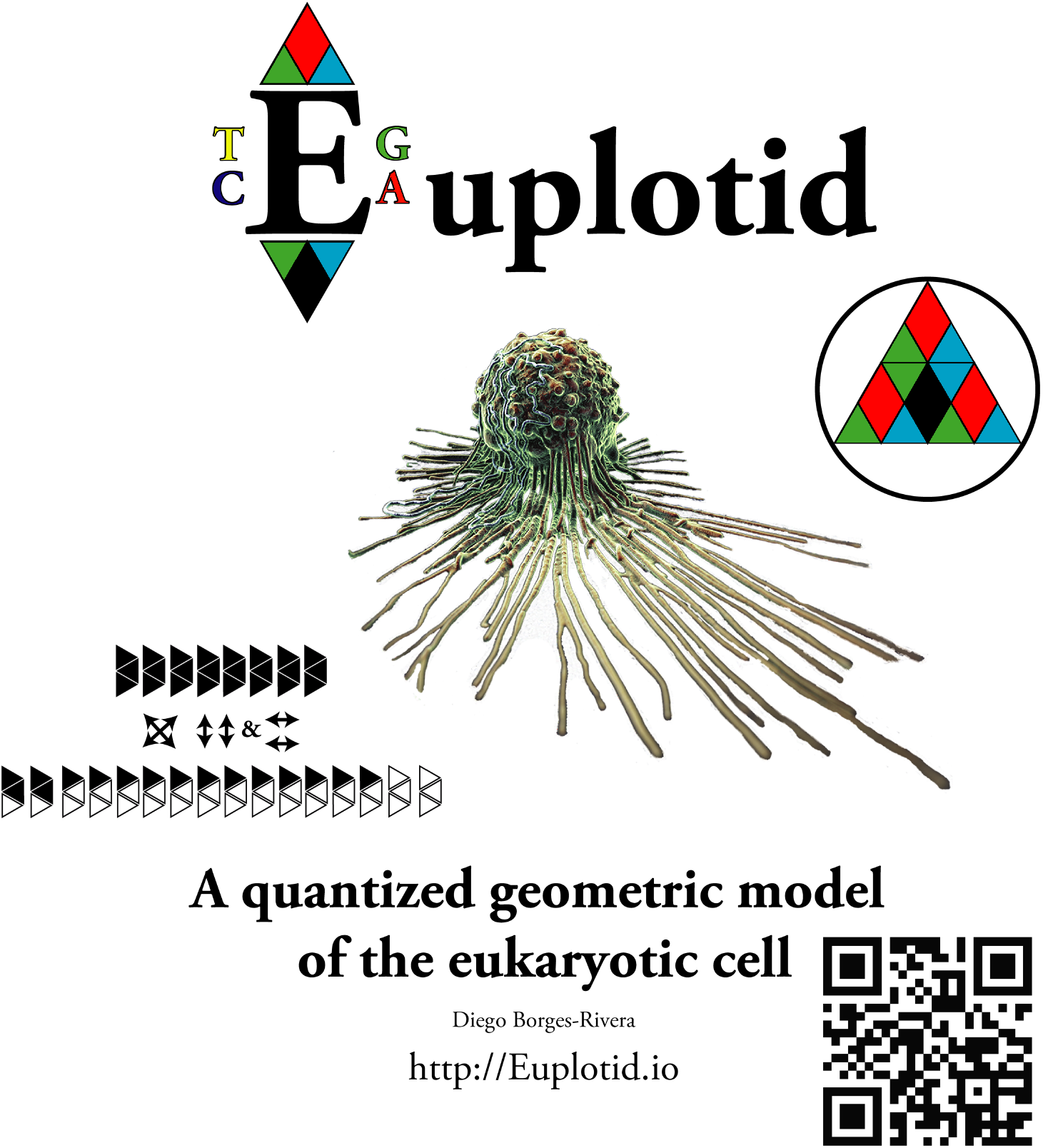

## 2 Epigraph: The Naming of Euplotid

“The cell is a system that is capable of creating and conserving its own code” –Marcello Barbieri^1^

The Cambrian Explosion is responsible for the cementing of animal life on this planet; phylogenetic analysis appears to indicate that all metazoans originated from a common flagellated organism, something that likely resembled a Euplotid. Darwin has noted that this event does not seem to follow with a traditional evolutionary view, the speed and emergence of metazoans accelerated orders of magnitude when compared to the life that preceded them. Many hypothesis have been presented for the cause behind the Cambrian Explosion, with evidence coming from oxygen concentrations or critical evolutionary developments, but few have looked towards the genetic code for clues.

A number of patterns have been observed within the codons, but a particularly interesting combination of two rules allows for the writing of the Euplotid genetic code in a contracted manner. By combining Hisagawa-Minata’s^2^ grouping by codon redundancy and sorting by mass, with Rumer’s Bisection^3^ by transitions and transversions, and applying them to the Euplotid’s genetic code we are able to contract and arrange the codons as first shown in Makukov et al^4^. The process is described and carried out in Figure 1.1

**Figure 1.1:**
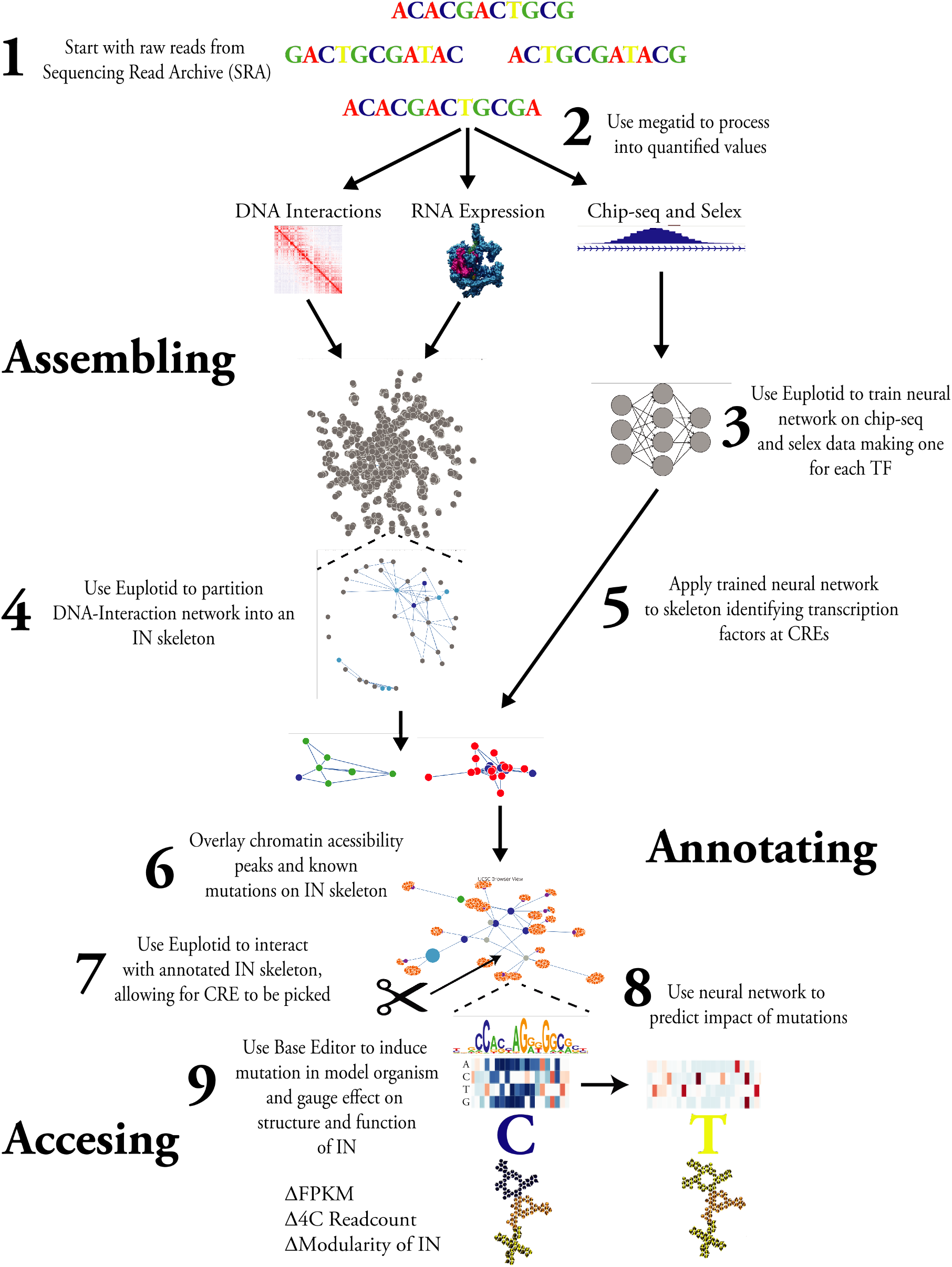
Detailed Abstract

Makukov et al discovered the protacted Euplotid codon arrangement and go well into the arithmetic interpretation. Figure 1.2 is an ideographical interpretation. Due to this unique genetic signature, I decided to name this quantized geometric model of the eukaryotic cell, “Euplotid”.

**Figure 2.1:**
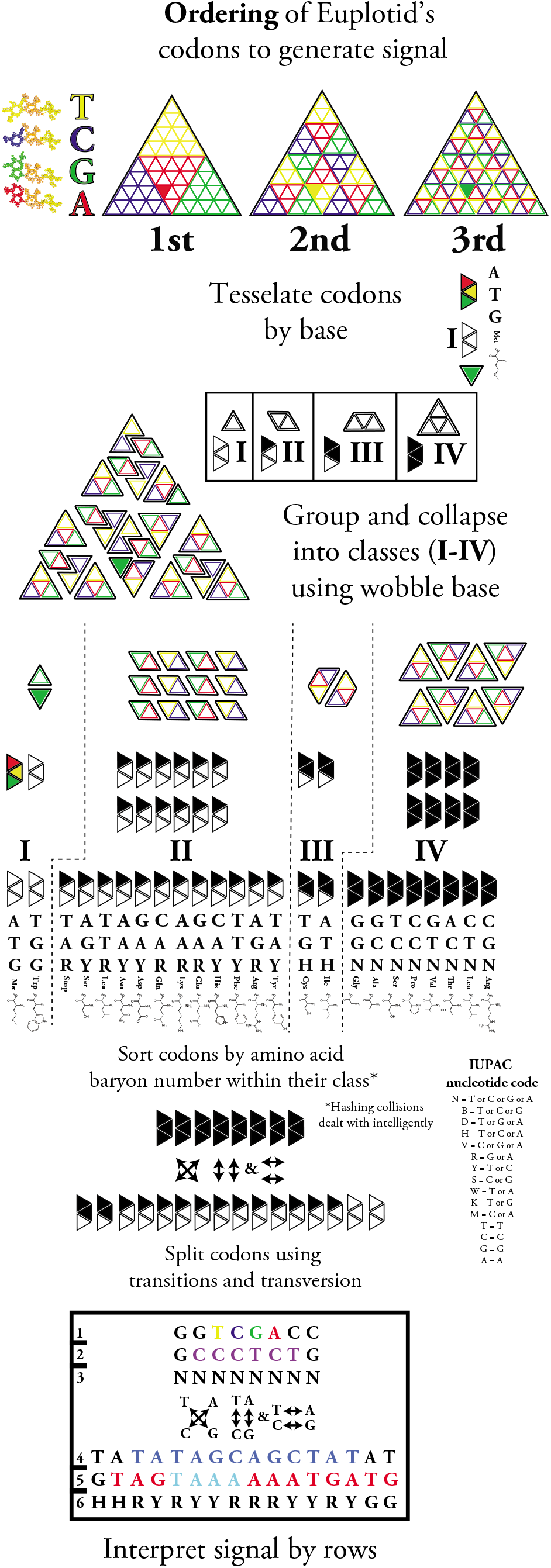
Ordering of Euplotid’s codons for generation of signal

**Figure 2.2:**
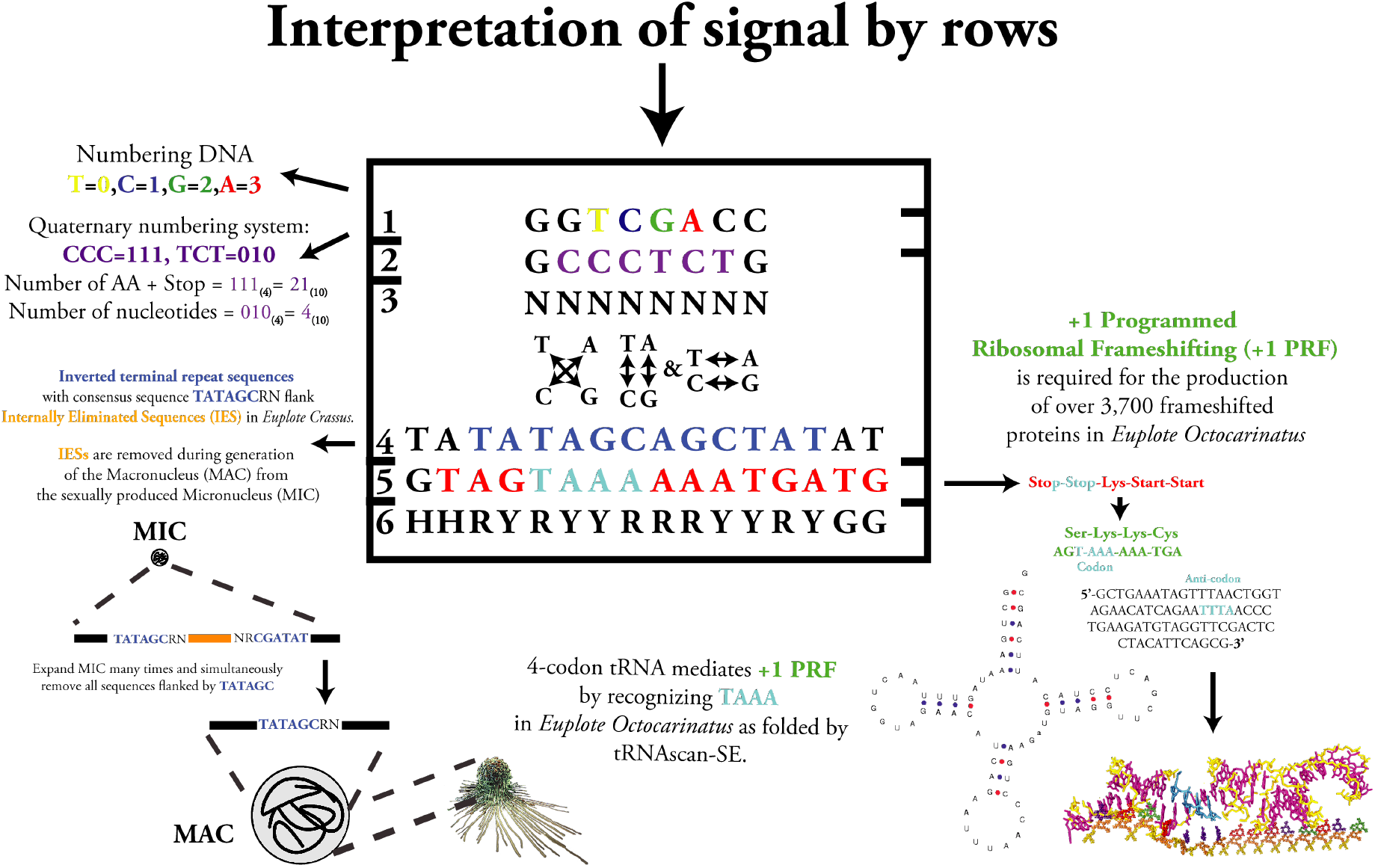
Epigraph - the naming of Euplotid:

1. GGTCGACC: How to number DNA: 0=T, C=1, G=2, A=3.
2. GCCCTCTG: What radix number system to use. CCC = 111 (base 4) = 21 (base 10) = number of amino acids + stop codon. TCT = 010 (base 4) = 4 (base 10) = number of nucleotides.
3. NNNNNNNN: Is able to complement “GGTCGACC”.
4. TATATAGCAGCTATAT: Euplotids remove Internal Eliminated Sequences (IESs) in the process of generating the large vegetative Macronucleus (MAC) from the small sexually produced Micronucleus (MIC). The Euplote Crassus consensus sequence is 5′-TATrGCRN-3′^5^.
5. GTAGTAAAAAATGATG: Translating from left to right starting w/ TAG would usually produce: Stop-Stop-Lys-Start-Start, but in Euplotids +1 Programmed Ribosomal Frameshifting (+1 PRF) occurs at AAA sites preceding stop codons due to a tRNA recognizing UAAA instead of UAA^6 7^. So if we perform a +1 PRF and read starting at AGT we get: Ser-Lys-Lys-Cys. +1 PRF in Euplotids is used to generate functional proteins from fusing different reading frames in over 3,700 proteins, including a number of Reverse Transcriptases^8^.
6. HHRYRYYRRRYYRYGG: Is able to complement “TATATAGCAGCTATAT”.

## 3 Introduction

The physical hierarchy of the genome begins with the smallest building block, planck’s quanta. The debate as to what geometric shapes define our 3D reality harps back to at least 360 B.C. when the philosophy behind the physicality of elements was established^9^ as exemplified in the Platonic solids. In Timeus some of the philosophical groundwork was laid for the understanding that compounds were made up of space-filling elements, and that through their combination one could create new compounds with very different properties than the sum of their parts. In the early 1800s the first atomic model was developed, able to explain chemical reactions as physical rearrangement of indivisible atoms^10^. The indivisibility of this atom was challenged in 1897 when the electron was discovered; the so called “plum pudding” model was born^11^. The plum pudding model was favored until 1911 when the infamous gold foil experiment proved that the positively charged “pudding” was actually a nucleus^12^. This nucleus contained protons, and later, was found to contain neutrons as well. Although the presence of the electron can be measured all around the positively charged nucleus they appear to be present at higher likelihoods in certain locations. During the 1930s a quantum electro dynamic model was born which is able to predict the probability of observing electrons at specific locations at unparalleled accuracy^13^. In tandem, particle physics gave us a clearer picture of the nucleus, as a tightly packed ball of protons and neutrons, each made up of quarks. This left us with the atom as a tightly packed nucleus surrounded by a field of electron “probability”, doomed to never truly know exactly what will happen. In 2012 an interesting proposition was put forth, what if a quanta of energy itself had a physical shape, albeit 2D? (fig 3.1)^14^. This gave rise to a unified way of interpreting all the previously developed theories, and coincidentally, loops back to Plato’s first solid, the tetrahedron (fig 3.2).

**Figure 3.1:**
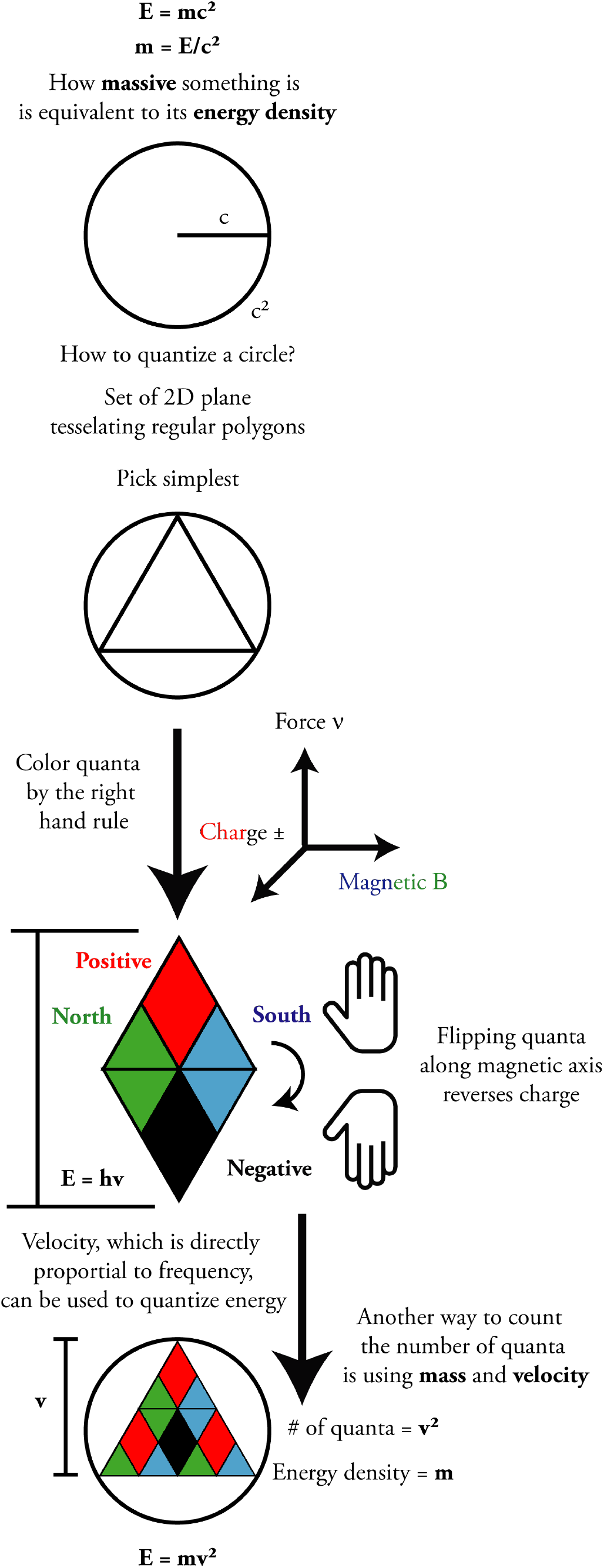
Quantizing an energetic plane.

**Figure 3.2:**
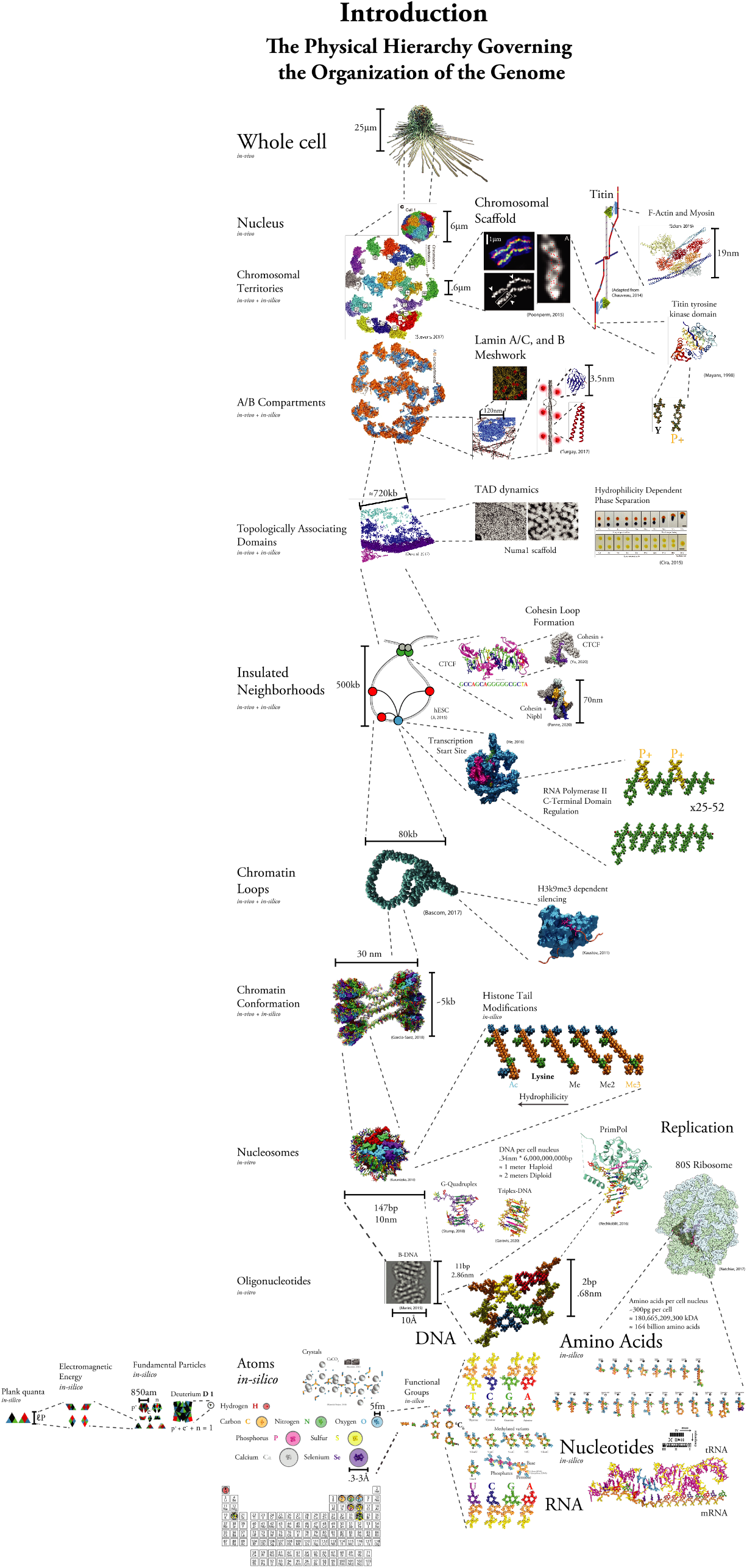
Introduction to the quantized 3D building blocks organizing the genome: We begin with planck’s quanta and use it to build the 3D blocks which comprise a eukaryotic cell. A brief video goes through the pseudo-physical model: [full introduction video](https://youtu.be/AflICQVeN44) while a 4D visualization is detailed at http://Euplotid.io.

Around 1860 the theory of natural evolution emerged describing how the diversity observed in organisms today may have come from a single common ancestor^15^. At the same time the physical explanation behind the tree of life was being laid down through the use P. Sativum^16^. The understanding that traits are inherited was extended to human disease in 1908 through the study of alkaptonuria^17^. Although the mechanism of inheritance was established, the physical material encoding the instructions for these traits was still hotly debated, was it proteins or nucleotides? The debate was settled in the 1930s through the use Pneumococcus and its virulence as a phenotypic trait^18^. With the chemical composition of the transforming material settled as nucleotides, the code defining the transition from DNA to RNA and then Amino Acids was solved during the 1950s^19^. The genetic code, which governs how an organism moves information between DNA, RNA, and Amino Acids, appears to be pliable and novel mechanisms continue to be found such as retrotransposable elements. Although we had found the chemical composition of the information storing component, we knew almost nothing as to how the order of DNA of is used to encode information. Two rules were discovered during the 1950s which began to decode DNA, Chargaff’s rules. The first rule laid the groundwork for the structure of DNA to be solved, that is to say %C=%G and %A=%T^20^. DNA’s structure was famously solved in 1953, giving a physical explanation to Chargaff’s first rule and a major step in the physical organization of the genome was solved, the 3D shape of an oligonucleotide^21^. The three prevalent double-stranded DNA structures were solved, A-DNA, B-DNA and Z-DNA, but other DNA structures continue to be discovered, recently evidence of DNA triplexes and G-quadruplexes (G4) has been recovered in-vivo, with their functions remaining enigmatic^22 23^. It is also very interesting to note that although a physical explanation was found for Chargaff’s first rule, his second remains unexplained.

Our understanding of the chemical composition of the genome was questioned when it was discovered that the canonical nucleotides existed within DNA in edited forms. The first conclusive proof of Cytosine Methylation on the 5th Carbon (5mC) was published in 1980^24^. In the following years a number of other modified states were found for Cytosine, including: 5-hydroxylmethylcytsine (5-hmC), 5-formylcytosine (5-fC) and 5-carboxylcytosine (5-caC), while the enzymes mediating them were found and characterized: TET, DNMT and AID^25^. Recently, evidence for mammalian DNA N6-methyladenine (N6-mA) has been recovered, although the recovery of the mark has proved to be extremely difficult, it appears to function in retrotransposon control^26^.

The first amino acid was discovered in 1806, asparagine. Over the next decade the rest of the canonical 20 amino acids were discovered, isolated, and their properties carefully measured^27^. The amino acids form the functional unit of peptides, which when strung together and folded, create proteins. Our understanding of how these mechanical subunits come together is still in its infancy, in much due to our misunderstanding of the charge mechanics at the most fundamental level. With a sharply defined boundary between energy and matter we are able to model interactions between these small protein building blocks in a far more natural, newtonian manner, while maintaining quantum accuracy.

Much of the confusion regarding the carrier of genetic information was due to DNA’s extremely tight association with positively charged protein complexes called nucleosomes^28^. The nucleosome’s components and structures were developed and refined throughout the 1940s and 50s, being refined to near atomic resolution. Although the predominant components were solved, new variants of the nucleosome complex were discovered, and continue to be, such as the newly characterized MacroH2A variants^29^. In the 1960s post-translational modifications of the nucleosome’s tail were discovered to affect its association with DNA through changing lysine’s charge and shape^30^. The nucleosome allows for about 200bp to be neatly packaged, tagged and accessed, laying the groundwork for information storage in larger diameter fibers.

Chromatin can be understood as any shape of DNA that has a diameter larger than the canonical 10nm beads on a string nucleosome model. The exact shape and the in-vivo existence of the 30nm chromatin fiber has been hotly debated. It appears that there exists evidence for both sides, and in reality, it seems likely that chromatin is a dynamic fiber, capable of deforming, memorizing, and reacting^31^. Within the last decade we have begun to probe how the shape and regulation of this fiber can impact its shape and function, we have begun to unravel the consequences of histone tail modifications on the conformation of chromatin. Chromatin is much more than the sum of its parts, and a key way of maintaining information in the shape of our genome^32^.

Once we had settled on DNA as the transforming material we could begin to elucidate the set of reactions which takes DNA and decodes it to amino acids through an RNA intermediate. Throughout the 1970s the first steps of RNA-Polymerase mediated transcription were solved and the key players identified^33^. It was quickly discovered that not all RNA is destined for translation, only the subset coined “messenger RNA” or mRNA. RNA-Pol II was shown to be the holoenzyme responsible for the polymerization of this mRNA. As the constituents of the pre-initiation complex (PIC) where discovered during the 1980s, the PIC was found to form around the canonical TATA DNA motif, coined the “TATA-box”^34^. This TATA box has a specific 3D shape which causes a bend of the DNA at approximately 80 degrees when bound by proteins called Transcription Factors (TFs). In Eukaryotes a protein “arm” complex called Mediator is attached to this TATA DNA bend acting as a transcriptional co-activator. Mediator, aided by Nipbl, allows for the threading of this bent DNA through a small proteinaceous band structure known as Cohesin, thereby creating a small loop^35^. The elongation of RNA-Pol II is preceded by TFIIH mediated phosphorylation of the C-Terminal Domain (CTD) of RNA-Pol II^36^. How the phosphorylation of the CTD impacts its interaction with the extremely electronegative surface of RNA-PolII has not been studied at the quantum level in the context of productive elongation.

The natural motor motion of RNA-Pol II during elongation serves to push the threaded Cohesin ring, causing an extruded loop, bringing linearly distant areas of the genome into close physical proximity. This extrusion process continues until a stopping block is encountered, such as a bound CCCTC-Binding Factor (CTCF) or a G4, stopping the progression of Cohesin and causing the ring to stack at that bound CTCF site. CTCF’s history in research took many turns, being assigned a plethora of roles, from enhancer to repressor, until its eventual establishment as a looping factor^37^. The rate of release of Cohesin at CTCF sites is also actively controlled by acetylation of the ring, while CTCF’s binding can be impacted by the methylation status of its DNA binding motif^38^. The intricate details are still debated, but overall it appears that the clever regulation of on/off rates on DNA for these three pieces, CTCF, Cohesin, and RNA-Pol II, allows for the creation of dynamic structures capable of reacting to differing cellular states, tuning a gene’s local regulatory architecture to adapt to specific environments. These structures have been coined Insulated Neighborhoods (INs) due to their ability of insulating CREs within the IN from those outside, definining CRE to TSS relationships (fig 3.3). Although a CRE is able to influence the expression of any gene in extreme genomic proximity, the larger structures encompassing the CRE can cause it to impact TSSs from seemingly distant promoters^39^. The local regulatory architecture of Eukaryotic genes is dependent on their own expression through the extrusion of Cohesin rings, this intriguingly forms a sort of feedback mechanism (fig 3.4). The speed and dynamics of this transcriptional feedback mechanism may be influenced by certain charge dynamics from localized areas of acetylation, such as those mediated by the H4 HAT complex or the asymmetrically loaded Acetylation of H3 at Super Enhancers^40^. Although these effects originate down from the very basic levels of the physical hierarchy, when aggregated together, may impact larger dynamics. We are beginning to see modeling approaches reaching the scales necessary to tackle chromatin looping questions, these models will continue to develop and gain in accuracy and generality. The physical hierarchy that controls the relation between CREs and their respective TSSs is a complex and extremely fine-tuned process, but it may be that this complexity originates from very simple building blocks.

**Figure 3.3:**
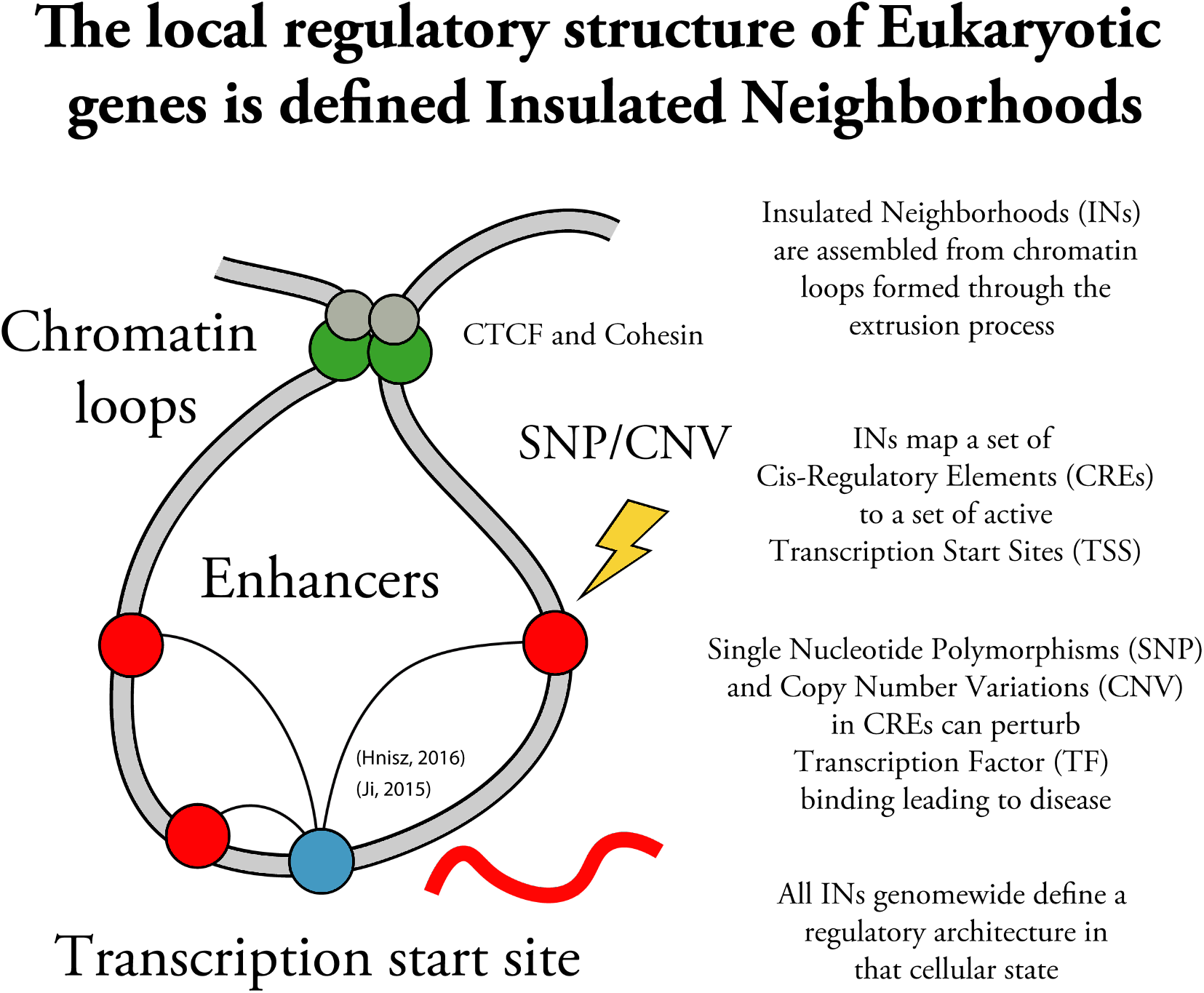
The local regulatory structure of Eukaryotic genes is largely maintained by clever use of CTCF and Cohesin whose misregulation can lead to disease.

**Figure 3.4:**
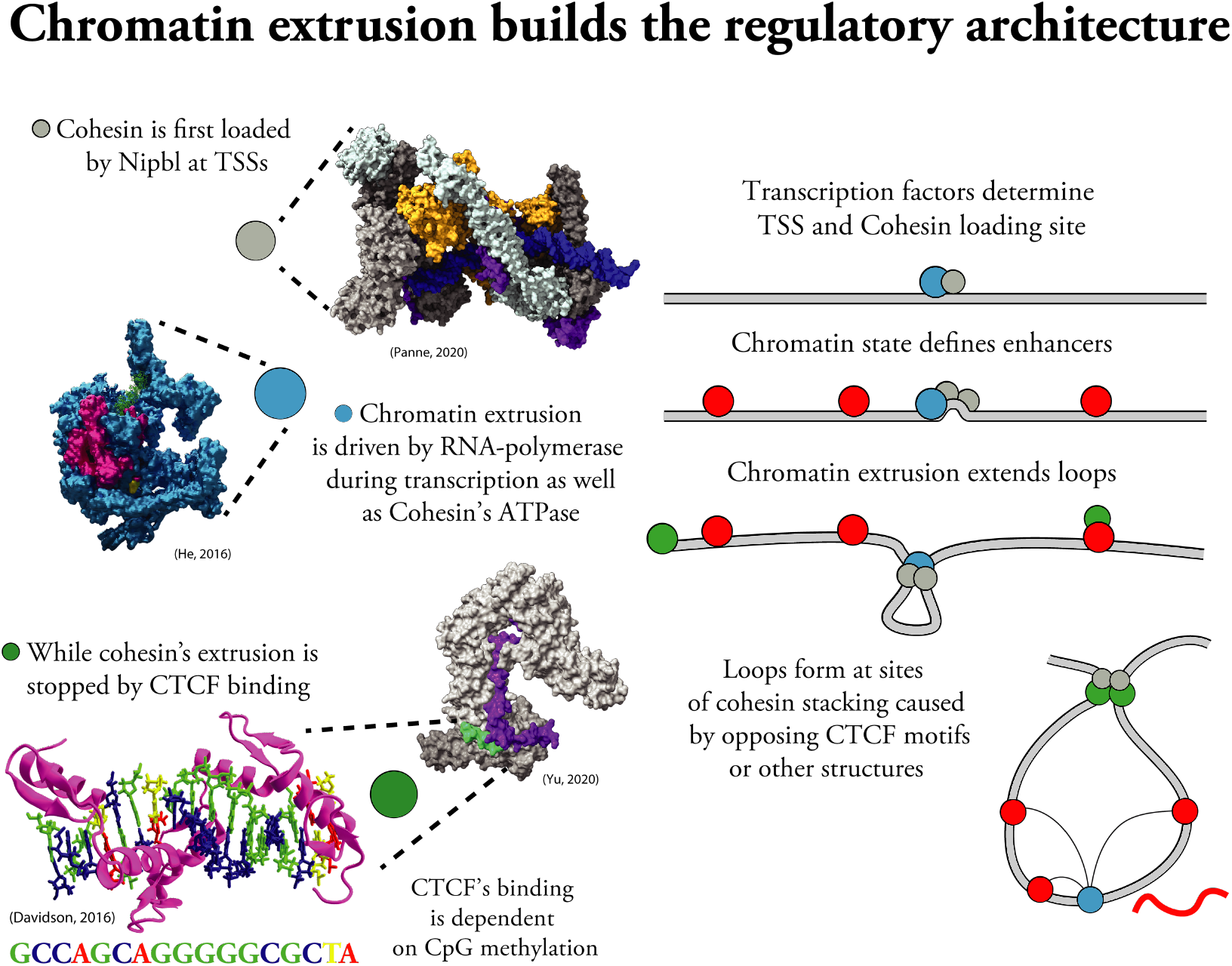
Chromatin extrusion allows for the stacking of Cohesin rings on CTCF

The regulation of the genome through the looping of CREs and TSSs appears to be often perturbed by disease causing mutations, especially those which are associated with non-coding CREs^41^, the vast majority of disease associated genomic loci fall within the 98% of the genome that is non-coding and therefore difficult to interpret (fig 3.5). By combining recently developed methodologies to probe RNA-Seq, DNA-DNA interactions and Chromatin Accessibility with our recently acquired knowledge of the physical hierarchy governing the folding of the genome, we are able to build a rough quantized picture of the 3D regulatory architecture of the genome. In order to digest this information in a manner which can guide experimentalists it is key to annotate and allow for easy access of these structures. Taking advantage of a number of recent developments in unrelated fields we are able to do just that; Euplotid provides a constantly evolving platform capable of allowing biologists to make physically informed decisions as to what variation is causative based on quantumly accurate models, virtually anywhere, anytime (fig 3.6).

**Figure 3.5:**
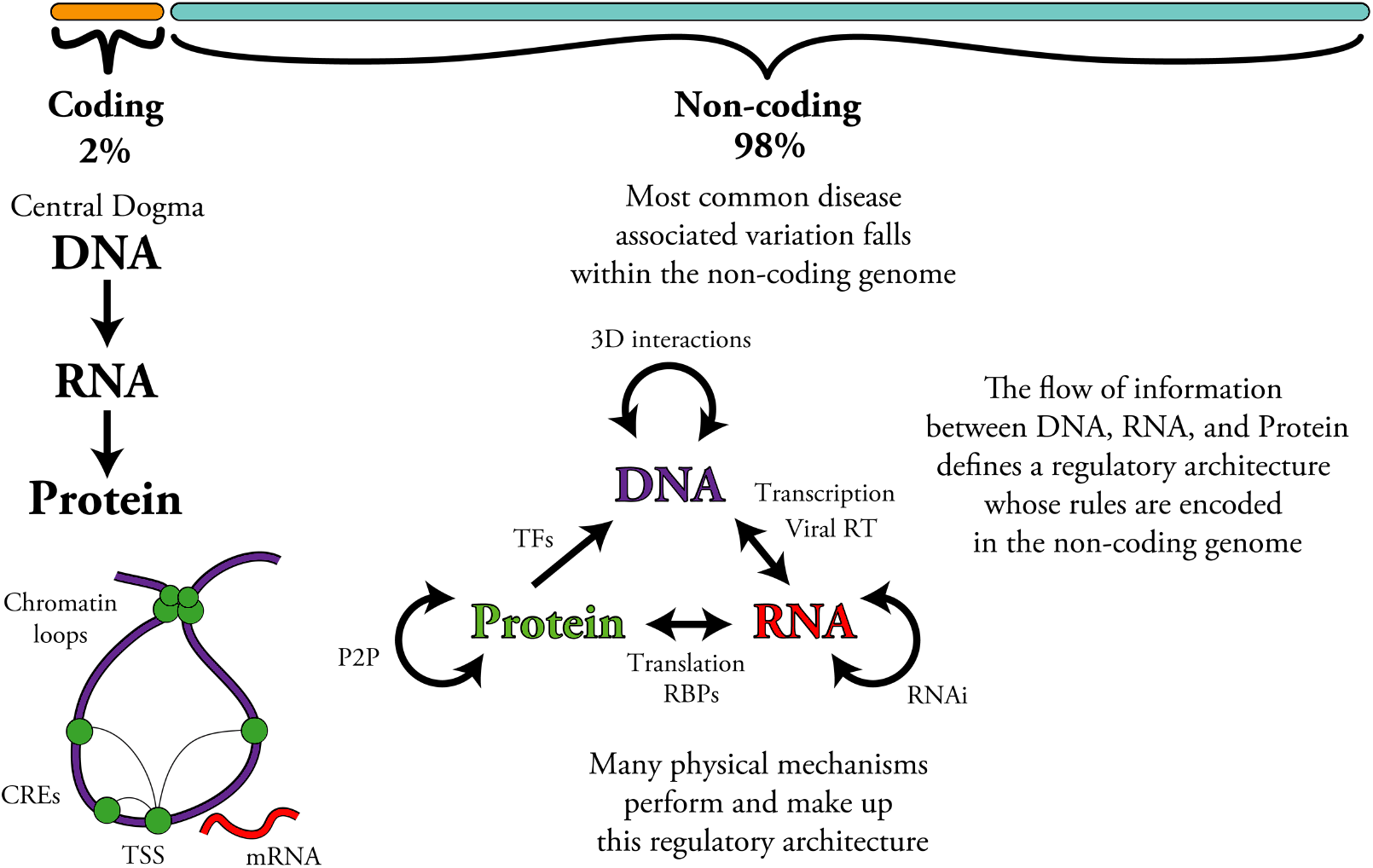
Euplotid problem

**Figure 3.6:**
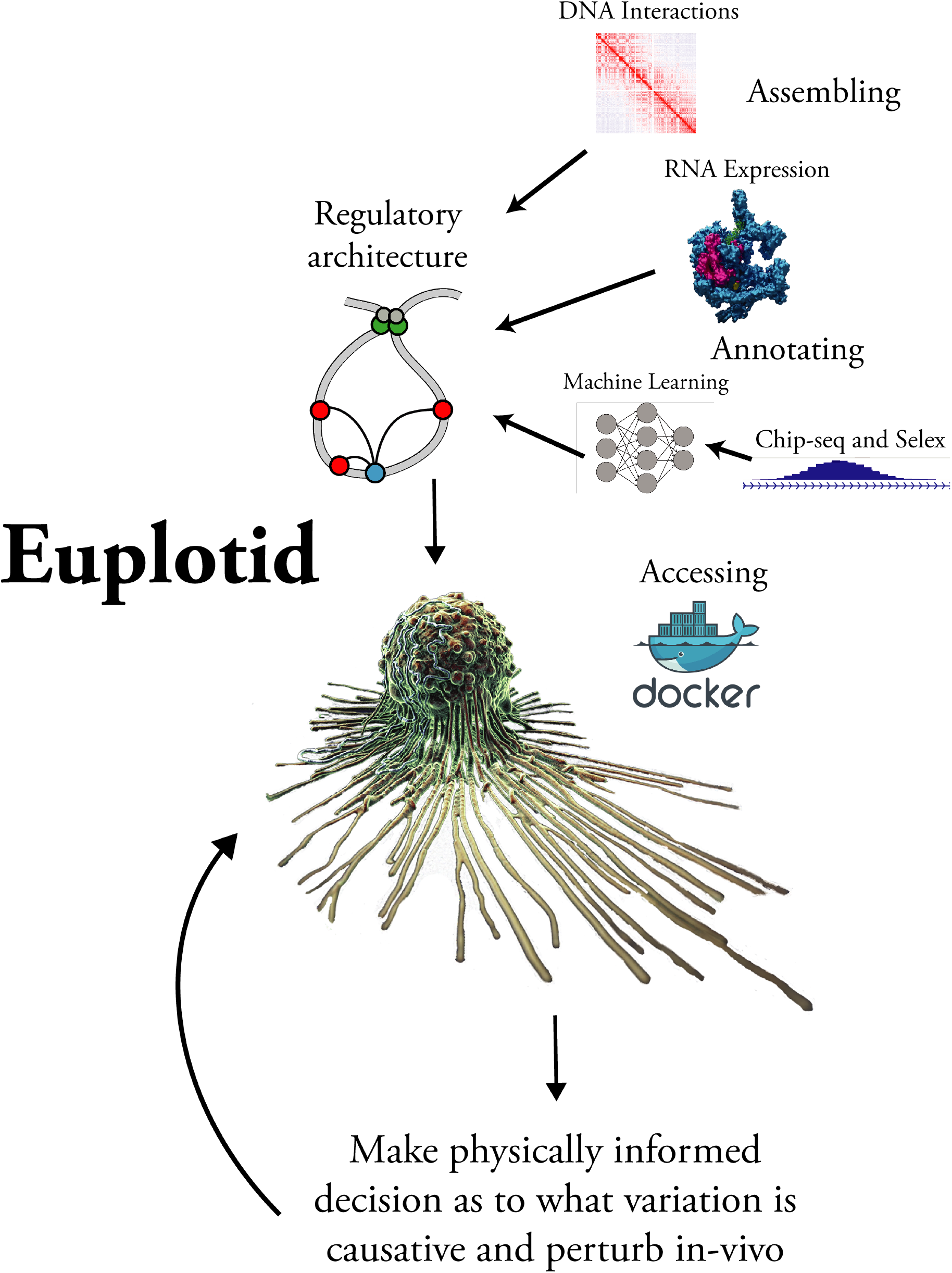
Euplotid solution

## 4 Results

### 4.1 Assembling

Determining the right local regulatory architecture of a gene can be easy by eye but defining and performing this task computationally is very difficult, here we present a louvain-based graph partitioning algorithm to chunk the DNA-DNA interaction of the genome into biologically relevant communities.

As shown in fig 4.1, we begin with a set of Nodes, defined as a DNA range, with a left and right boundary, for example: chr16:55155024-53806737. We then add a set of Edges, defined as DNA-DNA interactions, or loops. These loops can be recovered from living cells using a variety of methods, such as Hi-C, In-situ Hi-C, ChiA-PET, HiChIP, GAM, etc, each having their own wet and dry processing protocol, with dry protocols implemented within Megatid. During the starting condition (fig 4.1) every Node has its own community of size 1. We then use RNA-seq (RPKM) as a poor-man’s proxy for the rate of Cohesin-RNAPolII mediated chromatin extrusion. RNA-Seq reads were processed within Megatid using STAR to align the reads and RSEM to quantify RPKM^42 43^. We begin the algorithm by sorting the starting nodes by RPKM.

**Figure 4.1:**
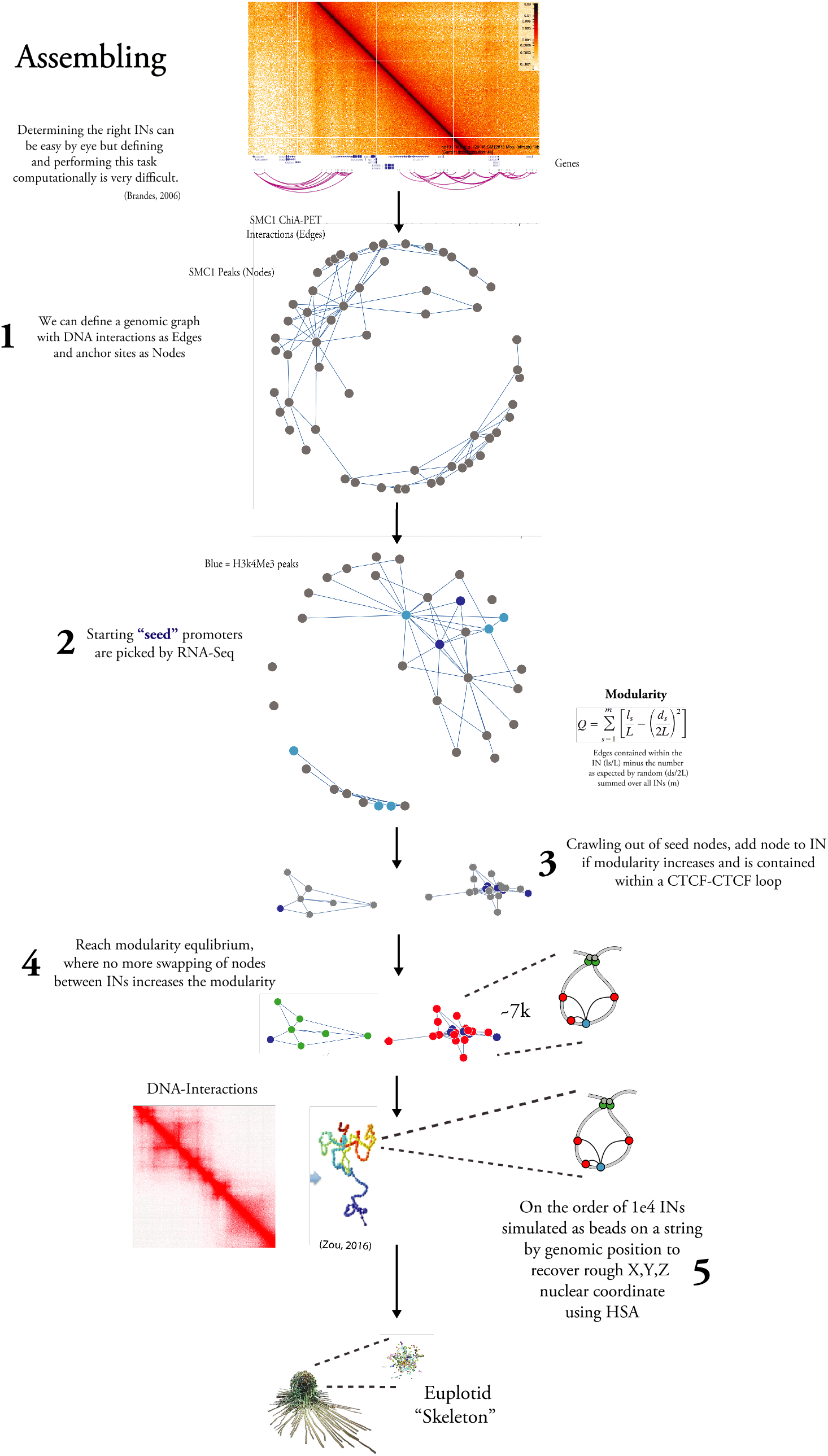
Assembling the local regulatory structure of genes

Beginning at each highly transcribed Transcription Start Site we then select the neighbor which would increase the overall network architecture the most as defined by Modularity and reassign its IN if the entire IN is encompassed within a single CTCF-CTCF interaction (fig 4.1)^44;45;46^. We then continue checking for valid reassignments for each node until no more moves exist which increase the overall graph’s modularity, thereby reaching a modularity equilibrium (fig 4.1). The nodes falling within one community are merged and then the process is repeated, creating hierarchical sets of communities. We then take all DNA-DNA interactions, and combined with Lamin A/B1 Chip-Seq, estimate the rough nuclear X,Y,Z position of each IN node by feeding the data through HSA^47^. The size of the node is defined using the sum of all reads falling within chromatin accessible regions (fig 4.1). This algorithm yields Euplotid’s skeleton, composed of about 10-15k INs.

### 4.2 Annotating

After assembling the Euplotid’s skeleton it is key to be able to visualize the impact of Histone modifications on the local regulatory architecture, therefore we color the nodes of the genomic graph by histone modifications in the given cell state of interest (fig 4.2). Specifically, red if the node overlaps only with H3K27Ac, blue if it overlaps H3k4me3^48^, and green if overlapping w/ CTCF.

**Figure 4.2:**
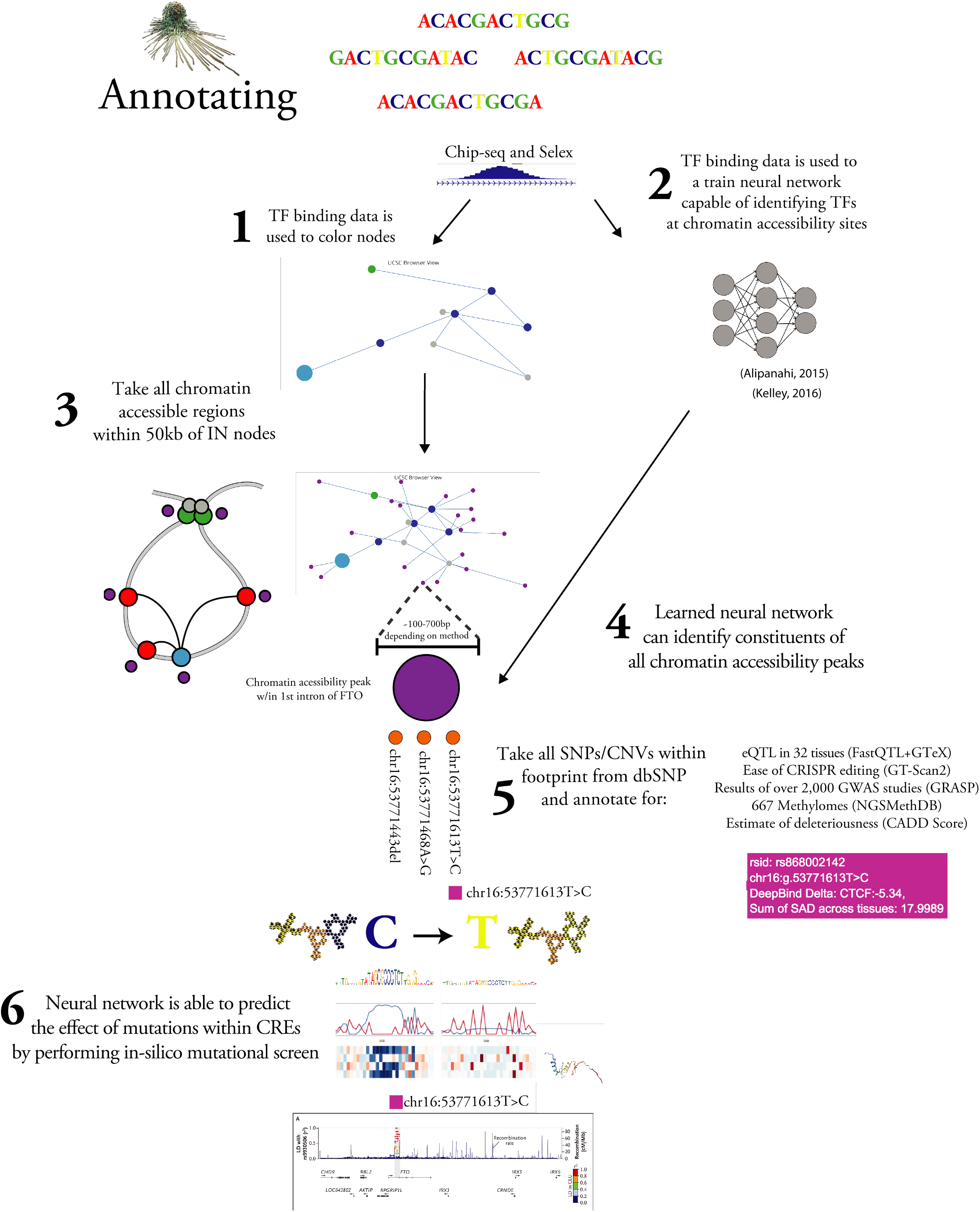
Annotating the Insulated Neighborhoods

Having a a good TF binding model allows for the identification of TFs within a given DNA; we begin by training Convolutional Neural Networks (CNNs) based on all chip-seq and SELEX data for all TFs surveyed using Chip-Seq or a similar enrichment protocol. The initial implementation of Euplotid uses pre-trained CNNs from Deepbind^49^. These CNNs are able to identify the TFs which fall under each chromatin accessiblity peak, but in order to understand the peak as a whole Euplotid takes advantage of Basset to train neural networks which are capable of predicting changes in chromatin accessibility^50^. Basset is trained on all available chromatin accessibility data in ENCODE, DNAse of 180 different cell lines. Basset is therefore able to perform in-silico simulations to gauge the impact of a given mutation on the the complex as a whole (SNP Accessibility Difference (SAD) profile), by combining this with the CNNs from Deepbind, we are able to make gleam insight as to what TF(s) are causing this change (fig 4.2).

We then take all chromatin accessibility peaks within a set distance (50kb) from all the nodes of a given IN allowing us to identify the relevant areas of chromatin which are actively being used in this particular cell state, some of which are acting as Cis-Regulatory Elements (fig 4.2). Any method of chromatin accessibility is appropriate, DNAse-seq, ATAC-seq, MNAse-seq, etc, all can be used as inputs. We then apply the trained neural networks on each chromatin accessibility peak, the activity of each TF neural network allows for the identification of a CRE’s constituents (fig 4.2). Currently this is performed by Deepbind, but a custom built pytorch based network will soon be implemented^51^ (fig 4.2). We can then include all SNPs/CNVs within dbSNP which overlap the chromatin accessibility peaks within the IN^52^ (fig 4.2). This variation is then annotated with the following data if available (eQTL in 32 tissues (FastQTL+GTeX), ease of CRISPR editing (GT-Scan2), results of over 2,000 GWAS studies (GRASP), 667 Methylomes (NGSMethDB), and an estimation of deleteriousness (CADD Score). For each variant which falls within the IN we perform an in-silico mutational analysis. This in-silico mutational analysis is as follows: predict the chromatin accessibility with and without the variant. The difference in SNP accessibility (SAD score) is calculated by Basset with the pre-trained networks as described above (fig 4.2).

### 4.3 Accessing

Visualization of large graphical structures with huge amounts of data which are navigable in a stable easy to deploy environment has been a huge barrier. Docker, Python, Resin, Plotly and Jupyter together allow for deployment, visualization, and interactivity across all computing architectures^53 54 55 56^. The entire analysis can be replicated and edited in front of your eyes, creating unprecedented transparency between input data and output analysis. Euplotid can be used to make mechanistic predictions from any device, in any web browser, running on virtually anything, from almost anywhere (fig 4.3).

**Figure 4.3:**
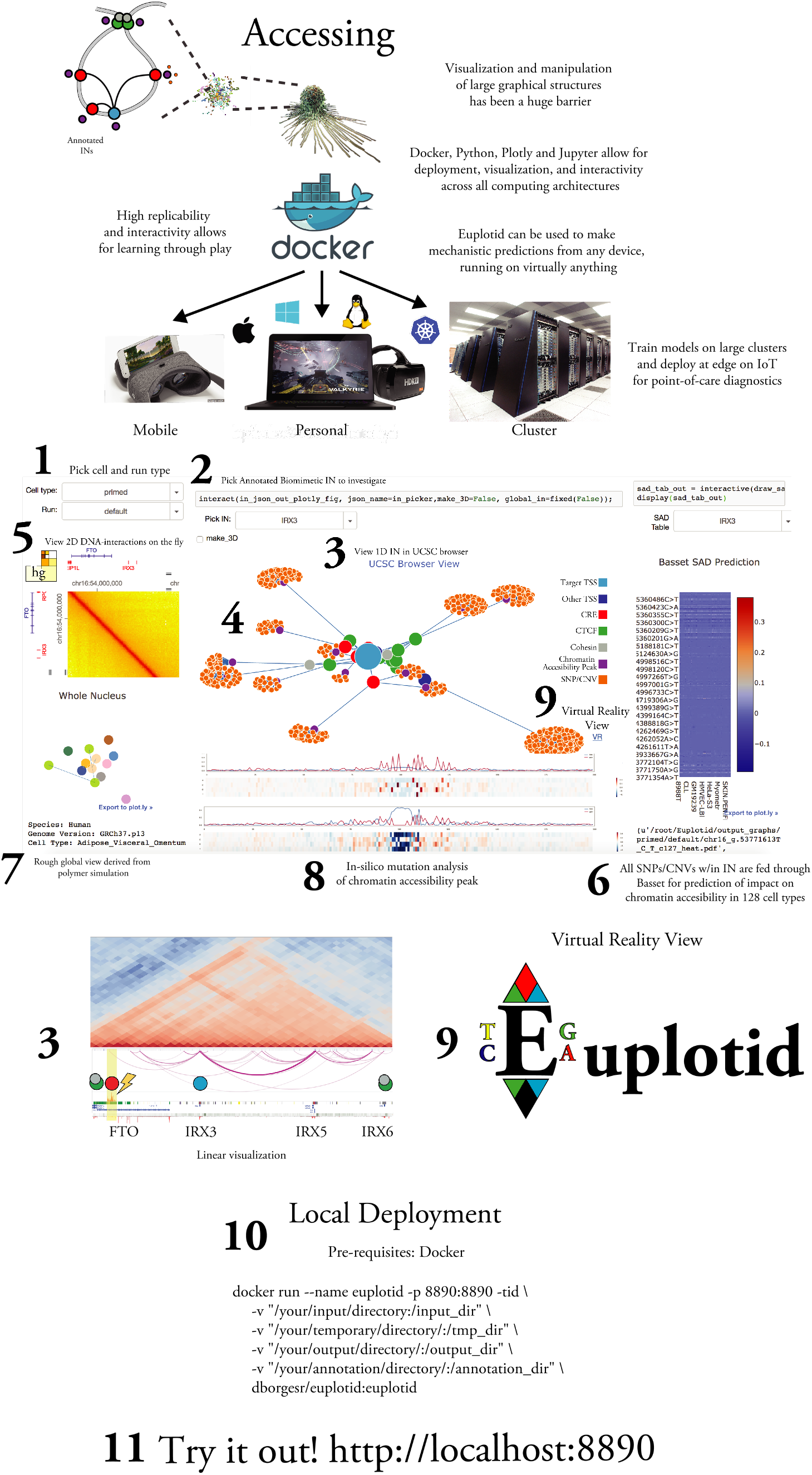
Accessing the Insulated Neighborhoods

Using Dash we are able to dynamically access the annotated local regulatory architecture of every gene which are stored as JSON graphs in the backend. Dash provides a simple way of interactively exploring data, a great way to explore the neighborhoods in 2D (fig 4.3). After picking a cell type and condition the list of annotated INs will be populated. A simple dropdown sorted by name is provided (fig 4.3). If the user wants visualize the data in the traditional 1D manner it is possible to load the data into a traditional genome browser, such as PyGenomeTracks^57^. To flatten INs to 1D we set the left and right boundaries as the leftmost and rightmost node within the IN currently being viewed (fig 4.3).

The annotated IN is positioned according to the Fruchterman-Reingold force-directed algorithm on DNA-interaction read count in order to have a more visually pleasing view. By employing 3Djs and Plotly we are able to navigate large graphical structures with relative ease. When hovering over every DNA-Interaction node the following pieces of data are shown if available: eQTL in 32 tissues (FastQTL+GTeX), ease of CRISPR editing (GT-Scan2), results of over 2,000 GWAS studies (GRASP), 667 Methylomes (NGSMethDB), estimate of deleteriousness (CADD Score), and CNN prediction of TF identity (fig 4.3).

Highglass is a tile-based viewing tool for DNA-DNA interaction data^58^. By employing Highglass we are able to quickly browse broad areas of the genome in a 2D manner, allowing us to confirm putative CRE to TSS relationships (fig 4.3). Employing CNNs previously trained on Chip-Seq and SELEX data and combining them with LSTM networks we are able to predict the chromatin accessibility of a particular sequence in a given cell type. Taking all SNPs/CNVs which fall within the IN we then predict the impact of each of those on the accessibility of chromatin (fig 4.3). Using the X,Y,Z coordinates generated for each IN using a simmulated annealing algorithm we are able to have a small “mini-map” corresponding to a global view of the entire nucleus. Each IN node is linked according to genomic coordinate while its coloring shows their chromosome (fig 4.3). Using previously trained neural networks we are able to view the predicted image for a given in-silico mutatation. This analysis gives an informed guess as to the impact a given SNP has on a Cis-Regulatory element, potentially affecting its function (fig 4.3).

Taking advantage of Virtual Reality (VR) technology developed for both the military and consumer markets we are able to render the annotated INs in full immersive VR. We use Unreal engine to design, build, and deploy the virtual world^59^. Deploying with the HTC Vive allows for fully immersive room-scale exploration of large complex annotated INs, while other platforms such as droid and iOS can join the same world in order to watch and interact. The 4D collaborative visualization can be downloaded for Windows, Mac, Android, and Linux at http://euplotid.io.

## 5 Discussion

Euplotid is a quantized geometric model of the eukaryotic cell that is built to evolve over time, to shed pieces and gain new ones as more powerful bioinformatic tools are created. Due to its modularity and deployability Euplotid can be used almost anywhere. Combined with emerging “Edge” computing and sequencing infrastructures such as NVIDIA’s Jetson and Oxford Nanopore’s MinION^60^ the potential for on-site building and visualization, from raw sequencing data to annotated immersive VR, will be possible.

In the future it will be possible to combine multiple images of Euplotid running in tandem mimicking tissues, with each Euplotid image communicating between each other and being slightly different, incorporating single-celled resolution techniques. Due to Euplotid’s foundational principles we are able to capture movement and mechanics down to the quantum level but remain extremely efficient and tractable, we render what we need to look at. The availability and ease of use will allow Euplotid to spread around the globe with relative ease.

## 6 Methods

The process of building Euplotid began with taking raw sequencing reads stored in a few different formats and processing it all the way to quantified values. Due to the pliability, breadth, and flexibility of the methods they will be documented within their own Jupyter notebook. Acting as both the documentation and the pipeline itself this format allows for seamless data integration.

- Hello world intro to programming and Jupyter’s capabilities [helloWorld](https://github.com/dborgesr/Euplotid/blob/master/pipelines/helloWorld.ipynb) O*.
- Databases and good tools to crawl the internet for interesting datasets and hypothesis [databases-Tools](https://github.com/dborgesr/Euplotid/blob/master/pipelines/databasesTools.ipynb) O*.
- Fetch any type of sequencing data from SRA [getFastqReads](https://github.com/dborgesr/Euplotid/blob/master/pipelines/getFastqReads.ipynb) O
- QC, trim, and filter sequencing reads fq2preppedReads O
- Call peaks from Chip-Seq and Chromatin Accessibility reads fq2peaks O
- Call normalized interactions from ChiA-PET reads fq2ChIAInts O
- Call normalized Interactions from HiC reads fq2HiCInts O
- Call normalized interactions from Hi-ChIP reads fq2HiChIPInts O
- Call normalized interactions from DNAse-HiC reads fq2DNAseHiCInts O
- Call normalized expression and counts from RNA-Seq reads fq2countsFPKM O
- Call differentially expressed genes from RNA-seq counts countsFPKM2DiffExp O
- Call normalized counts, miRNA promoters, and nascent transcripts from Gro-Seq reads fq2GroRPKM O
- Call normalized interactions from 4C fq24CInts O
- Build, annotate and add INs to global graph for a given cell state using DNA-DNA interactions, Chromatin Accesibility, and FPKM [addINs](https://github.com/dborgesr/Euplotid/blob/master/pipelines/addINs.ipynb) *
- View current built and annotated INs for all cell types [viewINs](https://github.com/dborgesr/Euplotid/blob/master/pipelines/viewINs.ipynb) O*.
- Search for and/or manipulate annotation and other data available to euplotid [annotationManagement](https://github.com/dborgesr/Euplotid/blob/master/pipelines/annotationManagement.ipynb) O*.
- Description of default software packages and images installed, how to get new ones, and which ones are currently installed. [packageManagement](https://github.com/dborgesr/Euplotid/blob/master/pipelines/packageManagement.ipynb) O*.
- Find clusters of interconnected nodes (Communities) using a Louvain algorithm then visualize the results [vanillaCommunities](https://github.com/dborgesr/Euplotid/blob/master/pipelines/vanillaCommunities.ipynb)
- Create, manipulate, and visualize cool DNA-DNA interaction files [chilledInteractions](https://github.com/dborgesr/Euplotid/blob/master/pipelines/chilledInteractions.ipynb)
- Design Base Editor and sgRNA plasmids for transition mutation at picked Cis-Regulatory Element CRE2plasmid

[O] = Megatid compatible [*] = Euplotid compatible [.] = Minitid compatible

